# Stage-specific transcriptomics of a leader cell reveals functional machineries driving collective invasion

**DOI:** 10.1101/2025.09.29.679269

**Authors:** Priti Agarwal, Iska Maimon, Hila Gingold, Victor Stolzenbach, Sarit Anava, Olga Antonova, Erin Cram, Oded Rechavi, Ronen Zaidel-Bar

## Abstract

Collective cell invasion underlies organ development, epithelial repair, and cancer metastasis. “Leader cells” remodel extracellular matrix, sense guidance cues, reorganize their cytoskeleton, and coordinate follower cells, but the molecular programs enabling these functions remain unclear. Here, we present a stage-specific transcriptomic dataset of the *C. elegans* gonadal leader cell, the distal tip cell (DTC), which invades basement membrane and guides germ cells to form U-shaped gonadal arms. Comparing invasive larval-stage DTCs with non-invasive adult-stage DTCs defines the molecular signature of an actively invading leader cell *in vivo*. Our dataset recapitulates known regulators of gonad morphogenesis and reveals numerous uncharacterized genes with potential roles in leader cell activity. As proof of concept, we identify vesicular trafficking proteins as enriched in invading DTCs, and demonstrate their importance for gonad development using endogenous tagging and DTC-specific RNAi. We also catalog diverse DTC-specific knockdown phenotypes. This resource establishes a molecular framework for leader cell activity and a platform to investigate conserved mechanisms of invasive migration.

## Introduction

Cell invasion, the ability of cells to migrate into adjacent or distant tissues, is fundamental to both normal development and pathological processes. Invasion occurs through two distinct modes: dispersed cell invasion (DCI), characterized by individual cell migration, and collective cell invasion (CCI), involving the coordinated movement of cohesive cell groups (Friedl et al., 2012; Prasanna et al., 2024; Wu et al., 2021). While DCI is exemplified by neural crest cell migration, CCI is central to numerous developmental events, including gastrulation, angiogenesis, and organogenesis, and is also a hallmark of solid tumor progression, often associated with higher metastatic efficiency (Ewald et al., 2008; Friedl and Gilmour, 2009; Jiang et al., 2013; Scarpa and Mayor, 2016).

CCI is a highly coordinated, multi-scale phenomenon that integrates extracellular matrix (ECM) degradation and remodeling, sustained mechanical coupling between cells, and supracellular coordination of signalling pathways (Mayor and Etienne-Manneville, 2016; Vilchez Mercedes et al., 2021). At the forefront of these invading cell groups are specialized “leader cells”, which drive invasion through ECM deposition, proteolysis, and remodeling. Leader cells also orchestrate cohesive movement by engaging in biochemical signalling and mechanical interactions with follower cells and determine invasion direction by sensing and responding to environmental guidance cues. Despite their pivotal role in CCI and tissue morphogenesis, the molecular and cellular mechanisms governing leader cell specification and their functions remain poorly defined, largely due to the difficulty of probing these dynamic interactions *in vivo*.

To address this, we employ *C. elegans* gonad morphogenesis as an *in vivo* model for studying CCI. This process involves invasion of a somatic leader cell, the Distal Tip Cell (DTC), followed by proliferating germ cells, recapitulating several aspects of collective cancer cell invasion. The *C. elegans* gonad consists of two symmetrical U-shaped arms encased within a basement membrane (BM). Each gonadal arm is capped by a single DTC, which also functions as a stem cell niche to promote germ cell proliferation behind it. Formation of the characteristic U-shaped gonad during the four larval stages depends on the directed invasion and guidance of the DTCs. The DTC actively remodels the ECM by secreting multiple matrix components and metalloproteases (Agarwal et al., 2022) and directs gonad morphogenesis by sensing chemotactic cues (Chan et al., 1996; Hedgecock et al., 1990; Leung-Hagesteijn et al., 1992) and selectively forming cell-matrix adhesions (Agarwal et al., 2022). The optical transparency and genetic tractability of *C. elegans* enables high-resolution live imaging and precise molecular perturbations, making the DTC a powerful model for dissecting the mechanisms underlying CCI and organogenesis *in vivo*.

Here, we define the stage-specific transcriptomic profiles of invasive (larval) and non-invasive (adult) DTCs to uncover the molecular programs driving leader cell invasion. Our analysis reveals invasion-specific gene expression signatures enriched for diverse cellular functions, including vesicular trafficking. Functional validation using DTC-specific RNAi (Linden et al., 2017) of candidate genes identified three novel vesicular trafficking genes – *sec-12, flwr-1*, and *arf-1* – as critical regulators of gonad morphogenesis. Additionally, we re-examined genes previously implicated in gonad morphogenesis to clarify their DTC-specific roles, revealing eight distinct phenotypic classes of gonadal morphogenesis defects. These findings highlight vesicular trafficking as a determinant of leader cell function and provide a valuable transcriptomic resource for investigating conserved mechanisms of collective invasion and organogenesis.

## Results and Discussion

### Stage-specific transcriptomic profiling of the distal tip cell

In larval stage 1 (L1), the gonadal primordium contains two germ cells and two somatic cells that give rise to the DTCs. During larval stage 2 (L2), the gonadal arms elongate along the ventral surface, each lead by a DTC. Proliferating germ cells provide the bulk force for elongation, while the DTC dynamically invades and remodels the surrounding BM through localized secretion of matrix metalloproteases. At larval stage 3 (L3), the DTC executes a turn away from the ventral surface towards the dorsal surface, by establishing polarized cell-matrix adhesions that generate torque and cause rotational movement (Agarwal et al., 2022). During larval stage 4 (L4), the DTC continues its invasive migration along the dorsal surface, extending the gonad towards the mid-body of the worm to complete the U-shaped structure. In adulthood, invasion ceases, but the DTC remains essential as a stem cell niche, sustaining gametogenesis and gonad function (Figure 1A).

**Figure 1:**
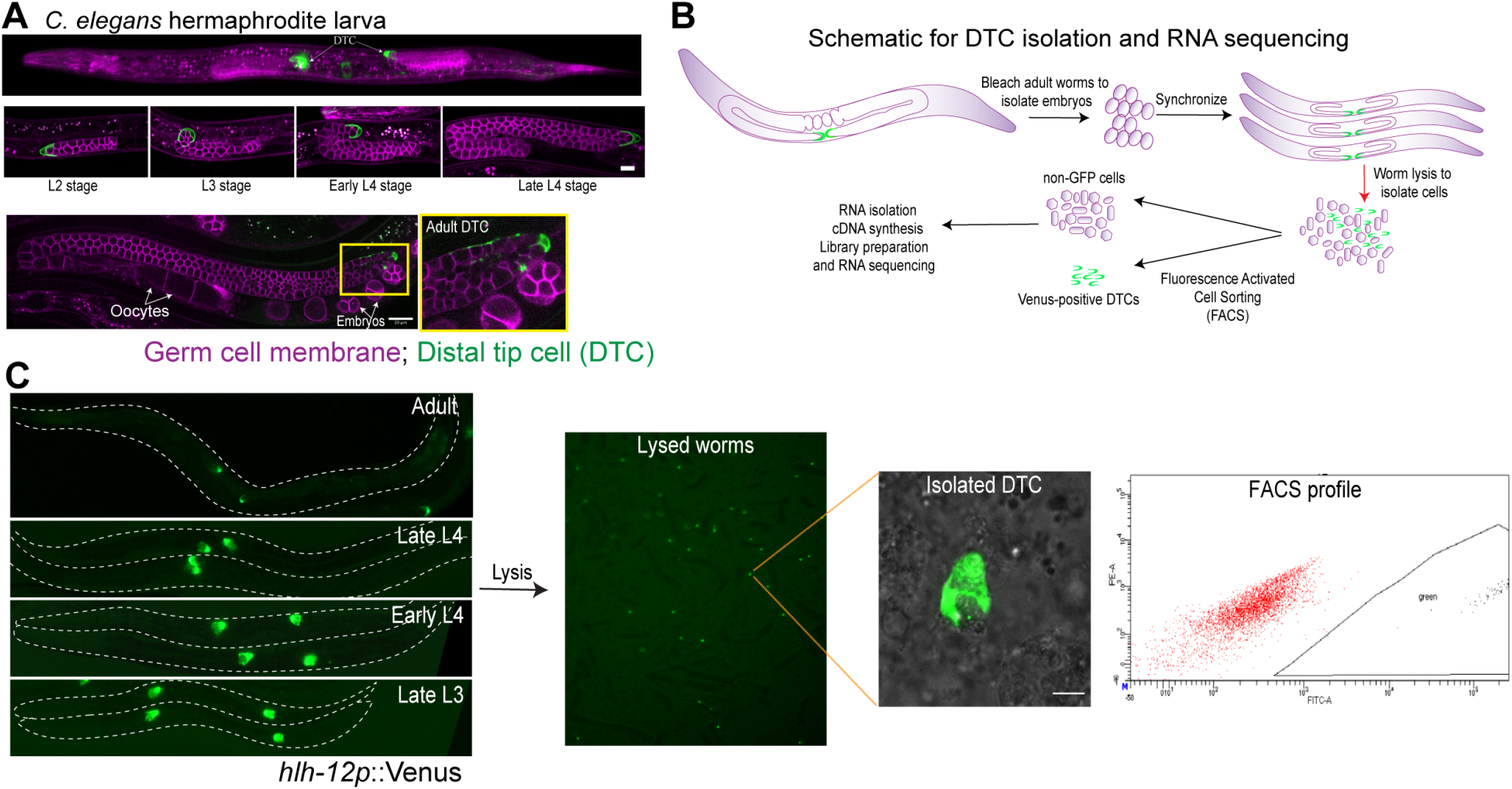
Stage-specific transcriptomic profiling of the distal tip cell. **(A)** Representative confocal images of gonads expressing a germ cell membrane marker (*pie-1p*::mCherry::PH(PLC1delta1, magenta) and a DTC membrane marker (*lag-2p*::mNG::PH, green) during indicated larval stages and adult stage. **(B)** Schematic of DTC isolation protocol for RNA sequencing. **(C)** Representative images of an experiment following the protocol in (B) showing fluorescently tagged DTCs in worms at four different stages of gonad development, isolated DTCs and the FACS gate used for sorting.

To characterize the transcriptome of the DTC across developmental stages, we used a strain expressing the fluorescent marker Venus under the DTC-specific HLH-12 promoter. Fluorescently labelled DTCs were isolated from synchronized late L3, early L4 and late L4 larvae – stages when DTCs are actively invasive – and from adults, when invasion ceases. Cells were dissociated by mechanochemical disruption and sorted by fluorescence-activated cell sorting (FACS). For each stage, unlabelled cells collected during FACS served as negative controls (Figure 1B,C). RNA sequencing was performed on 24 samples: three biological replicates for each of the four stages, with Venus-positive DTCs and Venus-negative controls from each replicate.

### Validation of DTC transcriptome

Sequencing data showed ∼ 99% overlap with the *C. elegans* genome (Table S1). Gene expression profiles were highly correlated among biological replicates (Figure S1A and S1B). Consistent with accurate cell sorting, Venus transcript levels were strongly enriched in Venus-positive samples (Figure S1C). Principal component analysis (PCA) confirmed stage-specific clustering of DTC transcriptomes and clear separation from Venus-negative controls (Figure S2A). Expression levels of protein-coding genes in Venus-negative samples closely matched WormBase medians across multiple developmental stages, further validating accuracy (Figure S2B).

To assess potential germline contamination, we compared upregulated DTC transcripts to germline-enriched genes (Reinke et al., 2004) and found only 0.9-6.4% overlap (Figure 2A). To further validate sample specificity, we analysed tissue-specific marker genes: the DTC-specific marker *lag-2* exhibited robust enrichment in Venus-positive samples, while markers for hypodermis (*dpy-7*), spermatocytes (*him-3*), sheath cells (*lim-7*), muscle (*myo-3*), neurons (*rgef-1*), and intestine (*vha-6*) were consistently expressed at much lower levels (Figure 2B).

**Figure 2:**
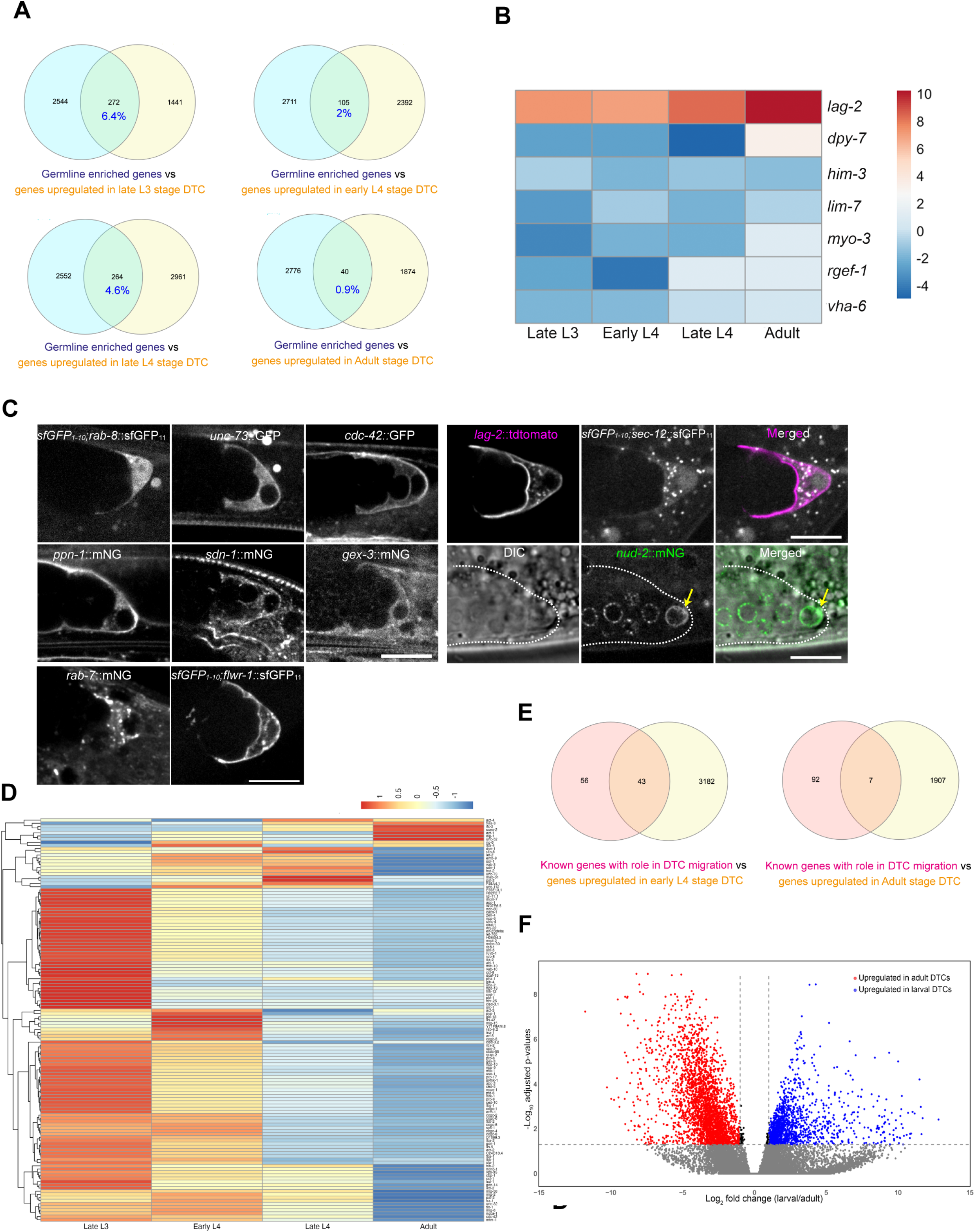
Validation of tissue specificity and stage-specific gene expression programs in DTCs. **(A)** Venn diagrams depicting the overlap between germline enriched genes (Reinke et al., 2004) and genes upregulated in late L3 stage DTC, early or late L4 stage DTC and Adult stage DTC. **(B)** Heatmap showing log_2_ fold change (L2FC) in expression of selected tissue-specific marker genes in DTCs compared to non-DTC controls at each developmental stage. Markers include genes associated with DTCs (*lag-2*), hypodermis (*dpy-7*), spermatocytes (him-3), sheath cells (*lim-7*), body wall muscle (*myo-3*), neurons (*rgef-1*), and intestine (*vha-6*).**(C)** Localization of endogenously tagged genes expressed differentially in the DTC transcriptome. **(D)** Scaled, clustered heatmap of expression for 99 genes previously implicated in gonad development (Cram et al., 2006), with expression values log-transformed, z-score normalized by row, and clustered using hierarchical clustering. **(E)** A Venn diagram depicting the overlap between the genes previously implicated in gonad morphogenesis (Cram et al., 2006) and genes upregulated in late L4 stage DTC and Adult stage DTC. **(F)** Volcano plot displaying differentially expressed genes between pooled larval stages (late L3, early L4, late L4) and adult DTCs, where genes with significantly higher expression in larval stages are shown in blue, those enriched in adults in red, and non-significant genes in gray; threshold lines indicate fold-change and significance cutoffs.

To corroborate the transcriptome, we examined protein localization for eight differentially expressed genes not previously reported to localize within the DTC: *unc-73* (a RhoGEF), *cdc-42* (a RhoGTPase), *ppn-1* (an extracellular matrix protein, papilin), *sdn-1* (a transmembrane proteoglycan), *gex-3* (a WAVE/SCAR complex component), *rab-8* (a Ras GTPase), *rab-7* (a Rab GTPase), and *nud-2* (a nuclear envelope protein). These genes were selected because endogenously tagged strains were available from the CGC. High-resolution live imaging confirmed DTC expression of all eight proteins with distinct subcellular patterns: RAB-7 in puncta, NUD-2 at the nuclear membrane, and the remaining distributed uniformly within the DTC (Figure 2C). Together, these results confirm both the accuracy and tissue-specificity of our DTC transcriptome.

### Stage-specific gene expression programs in DTCs

Differential expression analysis with EdgeR compared pooled larval stages (late L3, early L4, late L4) and adult DTCs, focusing on 99 genes previously implicated in DTC migration (Cram et al., 2006). Most of these genes peaked in late L3 and early L4, coinciding with active migration, while few reached maximal expression in adulthood (Figure 2D). More precisely, 43 genes were enriched in invasive early L4 DTCs and only 7 in non-invasive adult DTCs (Figure 2E). Genes not enriched in our dataset despite known functions in gonad morphogenesis probably act in other tissues and not in the DTC.

A global larval-versus-adult comparison revealed widespread transcriptional reprogramming: larval-stage DTCs upregulated 1,700 to 3,000 genes and downregulated 2,700 to 3,800 genes relative to other cell types, while adult DTCs upregulated ∼1,900 genes and downregulated ∼1,200 genes compared to non-DTC cells (Figure S3). Volcano plot visualization highlighted larva-enriched (red dots) and adult-enriched (blue dots) transcripts (Figure 2F).

### DTC-specific RNAi screen identifies regulators of gonad morphogenesis

To test whether the 43 genes enriched in larval DTCs act cell-autonomously to control gonad morphogenesis, we performed a DTC-specific RNAi screen followed by high-resolution phenotypic analysis. We used a DTC-specific RNAi hypersensitive strain (Linden et al., 2017) expressing germ cell and DTC membrane markers for live visualization of DTC morphology and gonad architecture (Figure 3A). Six genes with established DTC-specific functions (Agarwal et al., 2022) were excluded, leaving 37 candidates (Table S2). Of these, 17 produced no phenotype, while knockdown of 20 genes caused a spectrum of gonad morphogenesis defects, grouped into seven classes (Figures 3,4): (i) Fail to turn (Figure 3B), (ii) Fail to complete dorsal elongation (Figure 3C), (iii) Disorganized gonad architecture (Figure 3D), (iv) Duplicated gonadal arms (Figure 3E), (v) Gap between dorsal and ventral gonadal arms (Figure 3F), (vi) Pathfinding defects (Figure 4A), and (vii) Twisted gonads (Figure 4B).

**Figure 3:**
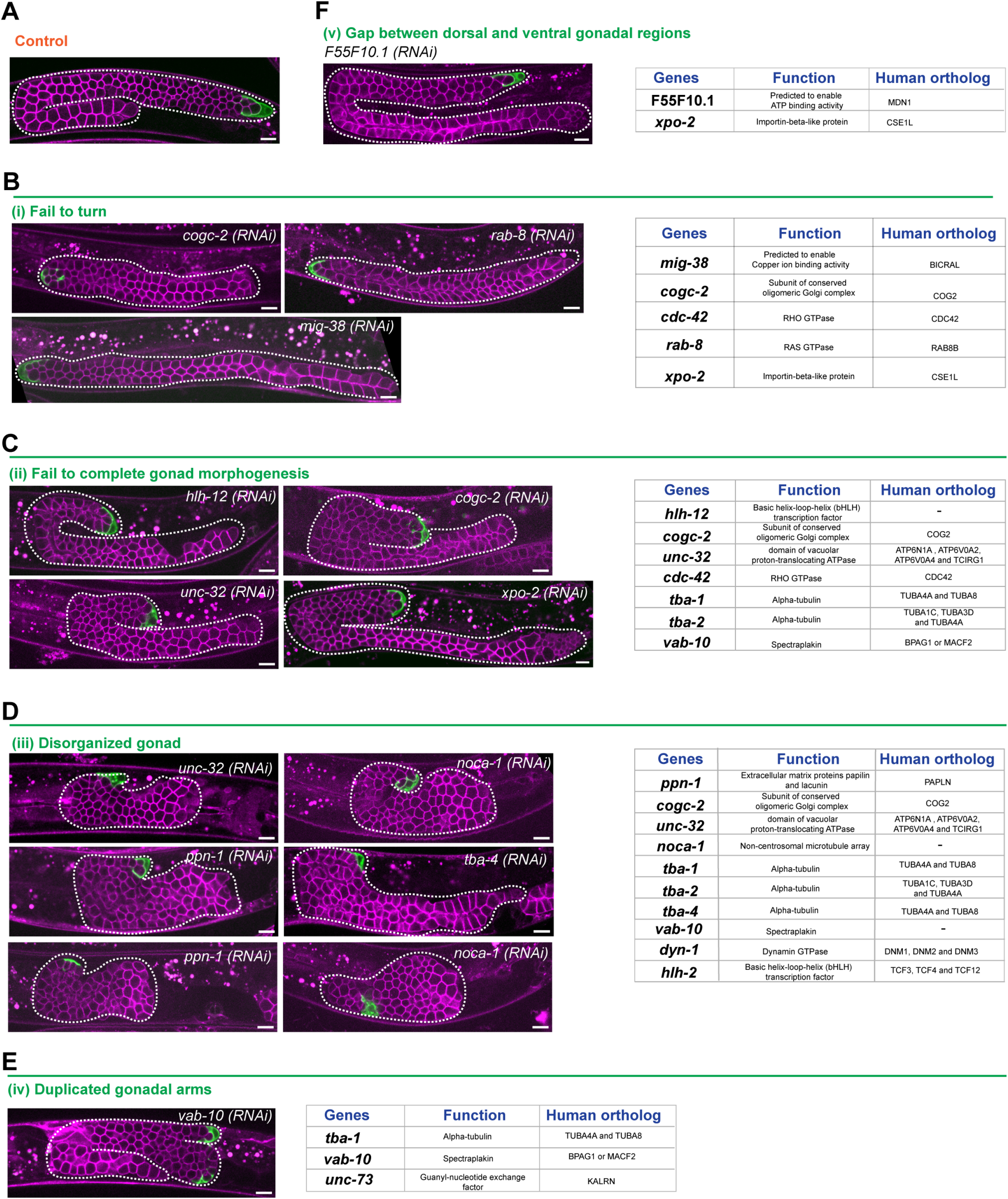
Gonad morphogenesis defects observed after DTC-specific knock-down. **(A)** Representative confocal image of a control gonad expressing a germ cell membrane marker (*pie-1p*::mCherry::PH(PLC1delta1), magenta) and a DTC membrane marker (*lag-2p*::mNG::PH, green). **(B-F)** Left: representative confocal images of the gonads expressing germ cell (magenta) and DTC (green) membrane markers after DTC-specific depletion of the genes indicated on the images. Right panel: list of genes exhibiting the same phenotype within a specific class, along with their functional classification and human orthologs, as reported in Wormbase. **(B)** Class (i) Fail to turn. **(C)** Class (ii) Fail to complete gonad morphogenesis. **(D)** Class (iii) Disorganized gonad. **(E)** Class (iv) Duplicated gondal arms. **(F)** Class (v) Gap between dorsal and ventral gonadal arms.

**Figure 4:**
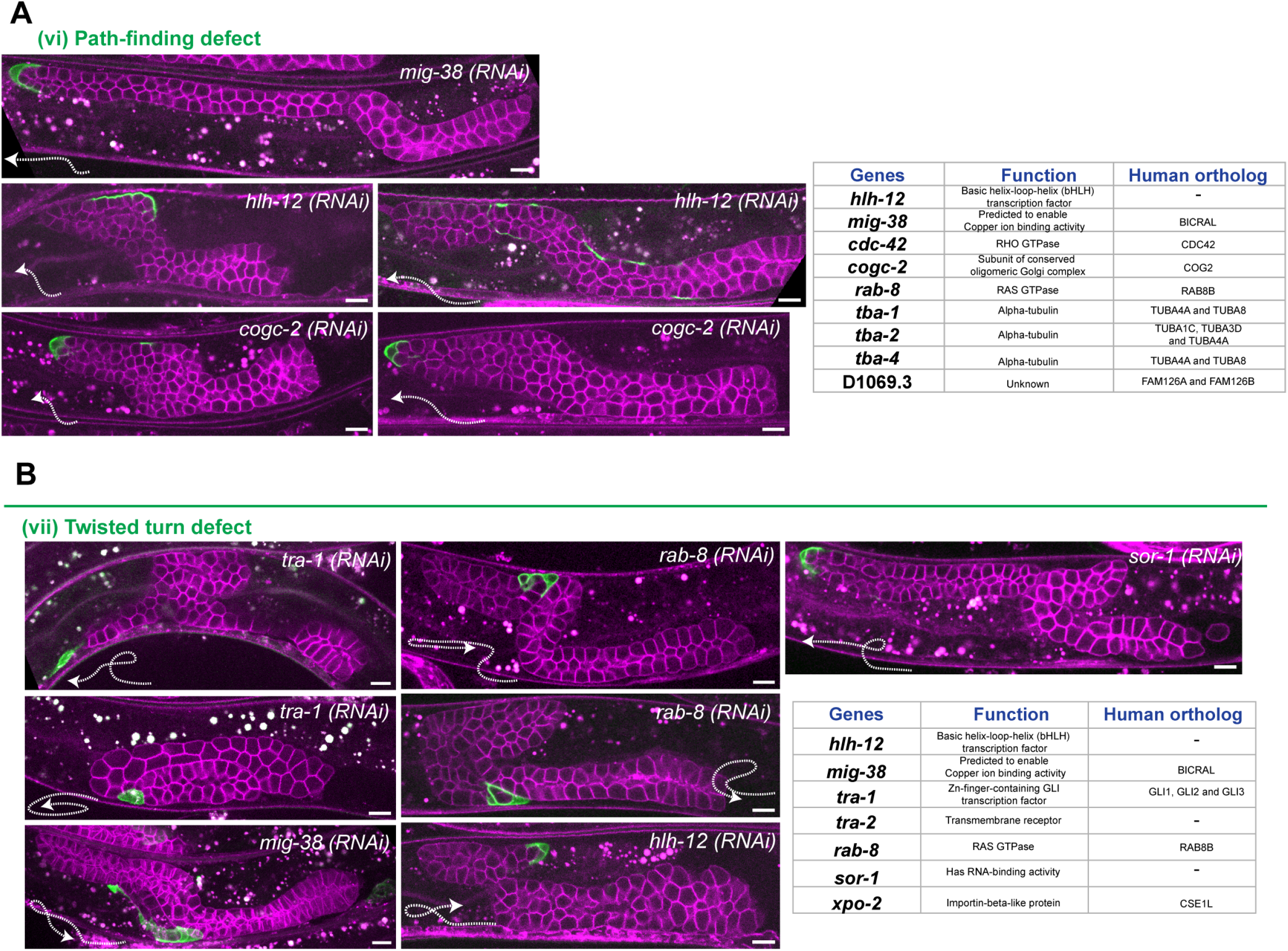
Turn defects observed after DTC-specific knock-down. **(A-B)** Left: Representative confocal images of gonads expressing a germ cell membrane marker (*pie-1p*::mCherry::PH(PLC1delta1), magenta) and a DTC membrane marker (*lag-2p*::mNG::PH, green) after DTC-specific depletion of genes indicated on the image. Right: List of the genes showing the same phenotype in a particular class, their functional classification and human orthologs as indicated in the Wormbase. **(A)** Class (vi) Pathfinding defect. **(B)** Class (vii) Twisted turn defect.

Classes (i), (vi), and (vii) suggest disruptions in directional cues or invasive machinery, consistent with known roles of Netrin signaling (Chan et al., 1996; Hedgecock et al., 1990; Leung-Hagesteijn et al., 1992) and polarized cell-matrix adhesions (Agarwal et al., 2022). Hence, the defects observed in these categories could stem from disruptions in the perception of chemotactic gradients or the dynamic regulation of cell-ECM adhesions. By contrast, class (iii) likely reflects impaired ECM remodelling by the DTC, a process essential for gonad elongation through regulated matrix secretion and degradation (Agarwal et al., 2022; Blelloch and Kimble, 1999; Blelloch et al., 1999; Nishiwaki et al., 2000). Precisely how each gene within this category differentially impacts matrix remodeling and consequently manifests in distinct phenotypes warrants further investigation. Class (iv) phenocopies defects caused by the loss of LINC complex protein UNC-83 and non-muscle myosin-II, which disrupt nuclear positioning and contractility, leading to DTC fragmentation and subsequent gonad bifurcation (Agarwal et al., 2024; Singh et al., 2024). Therefore, it is plausible that new genes identified in Category (iv) may be involved in nuclear positioning and/or contractile force generation within the DTC. Finally, class (v) – a novel phenotype featuring a gap between gonadal arms – may arise due to interventions from surrounding somatic tissues or indicate a lack of cohesive interactions between the gonadal arms. Further investigation into this phenotype may provide insights into aspects of tissue cohesion or inter-tissue interactions during organogenesis.

Notably, genes within each phenotypic class span diverse molecular functions, implying that multiple pathways converge within the DTC to regulate gonad morphogenesis. This observation challenges a previous proposal by Green and colleagues that shared gene knockdown phenotypes can be used to predict gene function (Green et al., 2011). In their study, similar gonad phenotypes were interpreted as indicative of shared molecular function or pathway involvement. However, we found that similar defects in gonad morphology can arise from disruptions in genes with very different molecular roles (Figure 4 and 5).

**Figure 5:**
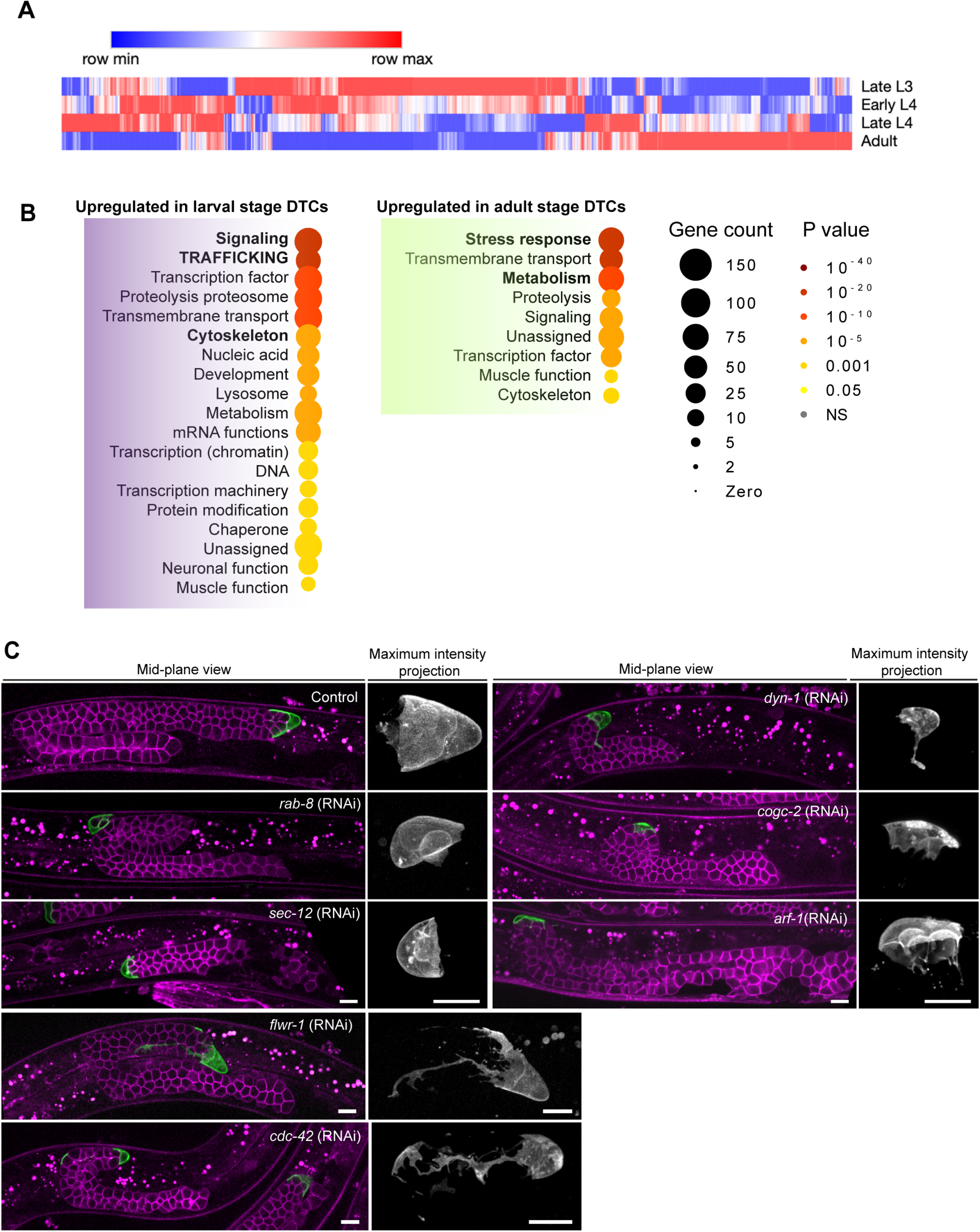
Trafficking genes are enriched specifically in the larval stage DTCs and affects gonad morphogenesis. **(A)** Global heatmap of gene expression in DTCs across four developmental stages: late L3, early L4, late L4, and adult. Genes with low expression were filtered out prior to plotting. Expression levels were scaled for each gene to highlight changes across developmental stages. **(B)** Gene ontology analysis of the differentially expressed genes in larval and adult stage DTCs (Holdorf et al., 2020) **(C)** Representative confocal image of gonads expressing a germ cell membrane marker (pie-1p::mCherry::PH(PLC1delta1), magenta) and a DTC membrane marker (lag-2p::mNG::PH, green) after DTC-specific depletion of trafficking machinery components indicated on the images.

Overall, the screen demonstrated that many genes previously linked to gonad development act cell-autonomously within the DTC, validating transcriptomic predictions and establishing a phenotypic atlas of DTC-specific loss-of-functions.

### Membrane trafficking genes regulate gonad morphogenesis

We performed global differential gene expression analysis using Morpheus. A heatmap of gene expression (Figure 5A) revealed two prominent gene clusters: one corresponding to late L3 and another to adulthood. Early and late L4 stages clustered closely together, exhibiting fewer uniquely expressed genes. Notably, early L4 genes overlapped extensively with those enriched in late L3, reflecting continuity in the molecular events driving active migration. By contrast, late L4 genes showed limited overlap with adult-stage genes, suggesting the onset of transcriptional changes marking the transition toward non-invasive stage.

Gene set enrichment analysis revealed that larval DTCs were enriched for pathways associated with dynamic cellular processes, including signalling, vesicular trafficking, transcription regulation, and cytoskeletal organization. In contrast, adult DTCs showed enrichment in pathways associated with cellular maintenance, such as stress response and metabolic processes (Figure 5B).

Given our interest in the invasive function of DTCs during gonad morphogenesis, we focused on 68 genes significantly upregulated in larval stages and annotated as trafficking, signalling or transmembrane transport protein (Table S3). A two-stage RNAi screen identified three membrane trafficking components whose DTC-specific depletion disrupted gonad morphogenesis: SEC-12, ARF-1, and FLWR-1 (Figure 5C; Table S3).

SEC-12 depletion led to severely shortened gonads that failed to elongate or turn. DTCs were markedly reduced in size and accumulated vesicle-like structures not observed in controls (Figure 5C). As a COPII complex component, SEC-12 facilitates ER-to-Golgi protein transport (Barlowe and Schekman, 1993; Nakano et al., 1988). In *C. elegans*, depletion of SEC-12 impairs cuticle collagen secretion (Zhang et al., 2021), suggesting it may play a broader role in delivering ECM components or remodeling enzymes required for gonad morphogenesis. SEC-12 also mediates Wnt ligand secretion from the ER (Sun et al., 2017), raising the possibility that it may traffic chemotactic (UNC-5/UNC-40)(Hedgecock et al., 1990; Leung-Hagesteijn et al., 1992) or integrin receptors (Agarwal et al., 2022) crucial for DTC polarity and directional invasion.

Depletion of ARF-1 produced flattened DTCs (Figure 5C). ARF GTPases regulate membrane trafficking, actin cytoskeleton organization (Randazzo et al., 2000) and ER dynamics (Poteryaev et al., 2005). A previous study in mammalian cells and *C. elegans* has shown that retrograde transport of integrins is critical for cell migration and gonad morphogenesis (Shafaq-Zadah et al., 2016). Thus, ARF-1 likely influences DTC membrane/cytoskeletal remodeling to affect DTC shape. Alternatively, it could be an indirect effect: the premature termination of gonad elongation, possibly due to a defect in the trafficking of metalloproteases or integrins, might lead to increased pressure on the DTCs, causing them to flatten.

*flwr-1*(RNAi) led to elongated DTC projections and incomplete dorsal gonad extension (Figure 5C). FLWR-1 is homologous to human CACFD1 and *Drosophila* Flower protein (Yao et al., 2009). Flower function as a calcium channel and promotes endocytosis at neuromuscular junctions and at the immune synapse (Chang et al., 2018). Thus, FLWR-1 might be involved in calcium-dependant vesicle-trafficking events crucial for DTC morphogenesis. Given calcium’s role as a regulator of cytoskeletal dynamics (Tsai et al., 2015), FLWR-1 may also regulate actomyosin contractility. Consistent with this idea, the long DTC membrane extension observed after knockdown of FLWR-1 resemble loss of non-muscle myosin-II (Agarwal et al., 2024; Tolkin et al., 2024).

We confirmed the localization of SEC-12 and FLWR-1 within DTCs using endogenous split-sfGFP tagging (Goudeau et al., 2021). SEC-12 displayed a punctate distribution within the DTC, while FLWR-1 localized at the plasma membrane (Figure 2C). We also closely examined the phenotypes of other previously identified secretory pathway components, including RAB-8, DYN-1, COGC-2, and CDC-42 (Agarwal et al., 2022; Cram et al., 2006; Kubota et al., 2006), all of which showed distinct defects: *rab-8* (RNAi) worms had dome-shaped DTCs; *dyn-1*(RNAi) resulted in extremely small DTCs with accumulated vesicles; *cogc-2* (RNAi) DTCs were notably flattened while *cdc-42* (RNAi) worms had fragmented DTCs (Figure 5C).

Together, transcriptomics, DTC-specific RNAi, and endogenous protein labelling, identify membrane trafficking pathways as central regulators of DTC invasion during organogenesis. Our dataset provides a framework for dissecting how trafficking integrate with adhesion, chemotaxis, and cytoskeletal networks to orchestrate leader cell migration. Notably, in follow-up work, stage-specific collagen expression was shown to be essential for the cessation of DTC migration and gonad elongation (Stolzenbach et al., 2025). Thus, this resource delivers both mechanistic insight and a platform for future studies into leader-cell driven organogenesis.

## Methods

### *C. elegans* maintenance and strains used in the study

All strains used in this study were grown on the plates containing nematode growth media (NGM) seeded with OP50, an *E. coli* strain, at 20°C. For DTC isolation, we used NF2169 strain (*mig-24p::Venus*) provided by Kiyoji Nishiwaki’s lab (Kwansei Gakuin University, Nishinomiya, Japan). RNAi screen was carried out on DTC-specific RNAi hypersensitive strain expressing a germ cell membrane marker, RZB353 (*cpIs121 I; rrf-3(pk1426) II; rde-1(ne219)*V;ltIs44 [*pie-1p::mCherry::PH(PLC1delta1); unc-119(+)*] IV) (Agarwal et al., 2022). The genotypes of other strains used or generated for this study are enlisted in Table S4.

### RNA interference

RNAi experiments were conducted by using standard feeding protocol (Timmons, 2006). For RNAi, HT115 bacterial clones expressing dsRNA, against specific genes, were obtained either from Ahringer or Vidal RNAi library (Kamath and Ahringer, 2003). For some of the genes, RNAi clones were not available from either library. These genes were amplified from wild-type genomic DNA, cloned into T444T plasmid (Sturm et al., 2018), and then the plasmid was transformed into HT115 bacteria. Primers used for constructing RNAi clones are listed in Table S5. Overnight grown RNAi bacterial culture was diluted to 1:100 and grown again at 37°C until it reaches 0.5 OD. Then, the culture was induced with 1mM IPTG for 3 hours. After induction, the culture was spread on NGM plates containing 100μg/ml ampicillin and 1mM IPTG. Gravid adult worms were kept on the RNAi plates and allowed to lay embryos for 2 hours. Later, the adult worms were removed from the RNAi plates, and after 48 hours i.e., at the L4 stage, phenotype of the progenies was scored. HT115 bacteria containing empty L4440 plasmid (for the RNAi clones obtained from Ahringer or Vidal RNAi library) or empty T444T plasmid (for the RNAi clones obtained by cloning genes into T444T plasmid) was used as control.

### Isolation of DTCs and RNA extraction

DTCs were isolated using the protocol described previously for isolating neurons (Kaletsky et al., 2016). We used NF2169 strain (*mig-24p*::Venus) to isolate DTCs since it expresses Venus specifically in the DTCs. Worms synchronized at four different stages – Late L3 stage, early L4, late L4, and adult stage – were washed 5-6 times with M9 buffer to eliminate all the bacteria.The washed worm pellet (approximately 250μl) was transferred to Eppendorf tubes. Next, the worm pellet was rinsed with 500μl of the lysis buffer containing 20mM HEPES pH 8.0, 0.25% SDS, 200mM DTT, and 3% sucrose) followed by incubation with fresh lysis buffer for 6-7 minutes at room temperature. After the incubation in lysis buffer, worms were rinsed 5 times with M9 buffer and then incubated with 20mg/ml of freshly prepared Pronase (Sigma, P5147) for 15-20 minutes. During this incubation period, worms were further lysed by vigorous pipetting at intervals of every 2-3 minutes. The completion of worm lysis was examined by observing a small drop of lysed suspension on a slide and observing it under the stereomicroscope. Lysis was considered complete when no large worm fragments remained. Lysis was halted immediately by adding ice-cold 250μL of 2% FBS. The cell lysate was filtered through a 30μm filter (Pluristrainer) and collected into FACS tubes placed on ice. Venus-positive cells were FACS sorted at 4°C using BD FACS Aria II sorter into an Eppendorf tube containing 850μl Trizol solution (Thermo Fisher Scientific, Cat. No. 10296010). From each sample, non-Venus cells were collected in a separate Eppendorf tube as a control. For each stage, Venus-positive and Venus-negative samples were collected in triplicates. The miRNeasy Mini Kit (Qiagen, Cat no. 217004) was used for RNA extraction.

We obtained approximately 5-9 ng/μl of RNA from 30,000 to 60, 000 cells. RNA quality was tested using Agilent TapeStation and samples with RIN (RNA integrity number) value above 7 were used for cDNA synthesis.

### cDNA synthesis, library preparation and sequencing

We used SMART-Seq v4 Ultra Low Input RNA Kit (Cat. No. 634889) for cDNA synthesis. For each sample, 1 ng of RNA was used for cDNA synthesis. The quantity and quality of cDNA was determined using Agilent TapeStation. One ng of cDNA was used as input for preparation of sequencing libraries using the Nextera XT DNA Sequencing kit (Illumina). Libraries were prepared according to the manufacturer’s instructions. Quality and concentration were verified with TapeStation 2200 and libraries were pooled according to molarity for sequencing on the NextSeq 500.

### RNA sequencing analysis

mRNA libraries were first assessed for quality using the FastQC tool (Andrews, 2010) and were then aligned to ce11 version of the genome using bowtie 2 (Langmead and Salzberg, 2012). The aligned reads were counted using the python-based script HTSeq-count (Anders et al., 2015) and the Ensembl-provided gff file (release-94). Next, the samples were compared for differential expression using the R package DESeq2 (Love et al., 2014). Genes were regarded as differentially expressed if they pass the criterion of FDR < 0.1.

### Principal Component Analysis

Principal component analysis (PCA) was performed in R using the *DESeq2* package. Raw read counts were first normalized by estimating size factors within *DESeq2*. A variance stabilizing transformation (VST) was then applied to reduce mean–variance dependence and improve interpretability of the expression matrix. The transformed data were used to compute principal components with the plotPCA() function, and the resulting values were visualized with *ggplot2*. Samples were grouped according to developmental stage (GL2, GL3, GL4, GA) and non-green controls (NL2, NL3, NL4, NA), with distinct shapes and colors assigned to each stage for clarity. The analysis confirmed clustering of biological replicates and clear separation between developmental stages.

### Morpheus

For heatmap row ordering, we performed unsupervised hierarchical clustering on a normalized, log2-transformed expression matrix. Genes were first filtered to retain features with mean log2 CPM > 1 across all conditions. Pairwise similarities between gene expression profiles were computed using Pearson correlation, and converted to a distance metric as 1 − correlation. Hierarchical clustering was performed with average linkage (hclust, method = “average”) on the resulting distance matrix. The dendrogram order was applied to the expression matrix to generate a row-ordered table suitable for direct import into Morpheus for visualization.

### edgeR heatmaps

Differential expression analyses and heatmap visualizations were carried out in edgeR (R/Bioconductor). Raw read counts were filtered to remove non-gene features, normalized using the Trimmed Mean of M-values (TMM) method, and modeled with glmQLFit() to estimate dispersions and compute stage-specific contrasts between GFP-positive distal tip cells (GL2, GL3, GL4, GA) and their non-green controls (NL2, NL3, NL4, NA). Heatmaps were generated for two focused gene sets: (i) distal tip cell migration genes, using mean normalized expression values z-scored across stages, and (ii) tissue marker genes (dpy-7, him-3, lim-7, myo-3, rgef-1, vha-6, lag-2), using log2 fold changes from stage-specific contrasts. Heatmaps were produced in R with the pheatmap package and visualized in ggplot2.

### Fluorescence imaging

Live imaging was performed on a spinning disk confocal (Yokogawa CSU-W1) Nikon Ti2E microscope using a prime 95B sCMOS camera (Photometrics) and Metamorph software (Molecular devices, Sunnyvale, California). L4 stage worms were mounted on a 3% agarose pad placed on a glass slide with 5µl of 10 mM levamisole. To analyze gonad morphology, images were acquired using 60X oil-immersion Plan-Apochromat objective. To analyze DTC membrane morphology, images were acquired using 100X oil-immersion objective with Z-stack spacing of 0.4μm spanning the complete width of the DTC.

### Strain generation using CRISPR/Cas9-mediated genome editing

A previously established CRISPR/Cas9 approach (Paix et al., 2017) and Split-FP strains were used to endogenously label the RAB-8, SEC-12 and FLWR-1. Split-GFP_11_ (*sfGFP*_11_) fragment was inserted at the C-terminus of RAB-8 and FLWR-1 and N-terminus of SEC-12 in a strain expressing *sfGFP*_*1*–*10*_ in all the somatic tissues including DTC (CF4587 strain *muIs253[Peft-3::sfGFP*_*1*–*10*_*::unc-54 3’UTR Cbr-unc-119(+)] II; unc-119(ed3) III*) (Goudeau et al., 2021). CF4587 strain was injected with ribonucleoprotein complex [1μl of Cas9 enzyme (10 μg/μl), 0.5 µl of KCL (1M), 5μl of tracrRNA (0.4 μg/μl), 2 µl of targeted gene crRNA (0.6 mM) and 4.5 μl of targeted gene ssODN(1μg/μl)]. We included 1 μl of *dpy-10* crRNA (0.4 μg/μl) and 0.8 μl of *dpy-10* ssODN (1μg/μl) which creates a dominant mutation at *dpy-10* site) in the injection mix to identify roller worms after injection (Arribere et al., 2014). F1 heterozygous rollers were checked for fluorescence and F2 unmarked wild-type progeny was used for further study. Sequences of all the crRNAs, ssODNs and primers used for sequencing are as follows:

*dpy-10* crRNA: GCUACCAUAGGCACCACGAG

*dpy-10* ssODN: CACTTGAACTTCAATACGGCAAGATGAGAATGACTGGAAACCGTACCGCATGCGGTGCCTATG GTAGCGGAGCTTCACATGGCTTCAGACCAACAGCCTAT

*rab-8* crRNA: AGAAGAGCUUCUUCAGCAAC

*rab-8* ssODN:

GTGGATCGGGGACACAGAAGAAGAGCTTCTTCAGCAACTGGAGCTGCAATTTGCTTCGTGACC ACATGGTCCTTCATGAGTATGTAAATGCTGCTGGGATTACAGGAGGAGGATCCTAAATTTATT GCTCTCTCGCCATGTTCTGCCCTTT

Primers for sequencing: Forward primer*-* TTGAAGAACGCCGTGAAGTG,

Reverse primer-AATATGCGAGAGAGAGCGGAC

*sec-12* crRNA: CAGAUGACAAUCUUCGGCGG

*sec-12* ssODN: ATTTGCGTAAAAACTGTGTATTTATCACACAGATGCGTGACCACATGGTCCTTCATGAGTATGT AAATGCTGCTGGGATTACATCCTCCTCCATGACAATCTTCGGCGGTGGCCAATCGAAAAAAGC GCCACTTATCGGAG

Primers for sequencing: Forward primer*-* TGCAACAATGAGCTGCTTCG,

Reverse primer-GAGTCTGAAAAACCAGGCTAATGG

*flwr-1* crRNA: GAAUUUAUGGAAUGUUGGAU

*flwr-1* ssODN with two tandem repeats of *sfGFP*_*11*_: GGAGATCCGGCGTGGAGCCCACAGGTCAACCAATCCAACATTCCAGGAGGAGGATCCCGTGA CCACATGGTCCTTCATGAGTATGTAAATGCTGCTGGGATTACAGGAGGAGGATCCCGTGACCA CATGGTCCTTCATGAGTATGTAAATGCTGCTGGGATTACATAAATTCCTAAAAGTCATTTCTTC CCGCCCTCAAC

Primers for sequencing: Forward primer*-* GTTCGCTTCCTGGCAAAAGG,

Reverse primer-AGTTGGTCGTGGTCAAATTGC

## Supplemental material

**Figure S1:** Validation of DTC transcriptome

**Figure S2:** Stage-specific expression of DTC transcripts

**Figure S3:** Overlap of differentially expressed genes in DTCs between stages

**Table S1:** RNA sequencing alignment report

**Table S2:** DTC-specific RNAi screen of genes shared between larval-stage DTC transcriptome and known gonad morphogenesis regulators

**Table S3:** RNAi screen of trafficking-related genes enriched in Distal Tip cell

**Table S4:** The genotype of C. elegans strains used in this study.

**Table S5:** Oligos used for cloning of the genes of interest into T444T vector

## Data availability statement

All data are available in the main text or the supplementary materials. All NGS data are available through GEO, accession number GEO: [to be completed in revision].

## Acknowledgments

We thank Kiyoji Nishiwaki’s lab (Kwansei Gakuin University, Nishinomiya, Japan) for providing us the NF2169 strain (*mig-24p::Venus*). This research was supported by Grant No. 2021014 from the United States-Israel Binational Science Foundation (BSF).

## Author contributions

P.A. conceptualized the project, performed experiments, analyzed and visualized data, and wrote the paper; I.M. performed experiments; H.G. and V.S. carried out RNAseq analysis;

S.A. and O.A. helped with cDNA library preparation and sequencing experiments; E.C. and O.R. reviewed the paper, obtained funding, and supervised; R.Z.-B. contributed conceptualization, obtained funding, analyzed data, supervised, and wrote the paper.

## DECLARATION OF INTERESTS

The authors declare no competing interests.

## Notes

### Competing Interest Statement

The authors have declared no competing interest.

