## Supplementary Figures 1-3 for "Stage-specific transcriptomics of a leader cell reveals functional machineries driving collective invasion"

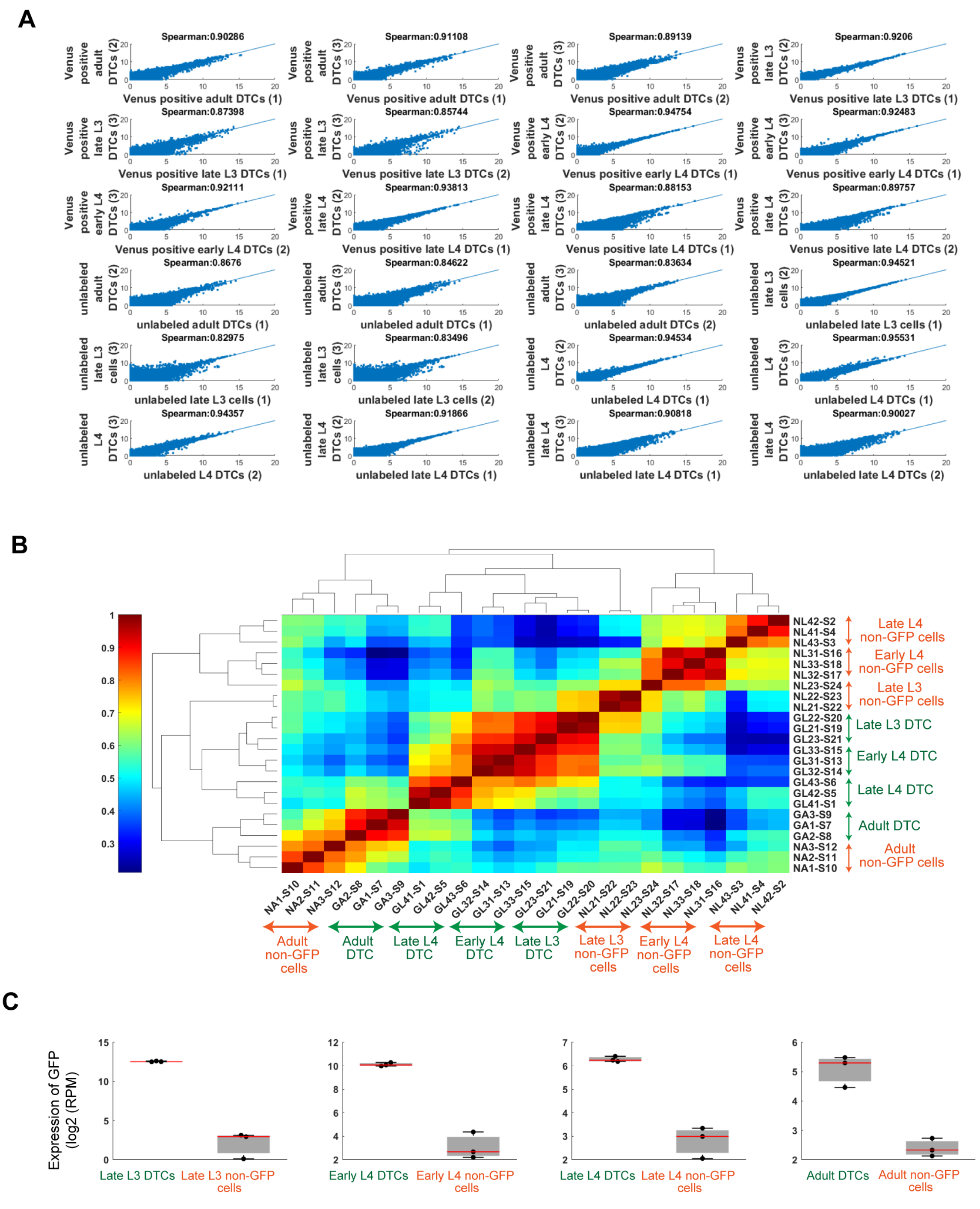


**Figure S1: Validation of DTC transcriptome.**
**(A)** Pairwise comparison of mRNA levels for protein-coding genes between replicate samples from the same developmental stage. Expression values are displayed as log2 (RPM + 1). **(B)** Hierarchical clustering of samples based on normalized mRNA levels of

protein coding genes. Off-diagonal entries show Spearman correlation between pairs of samples. The analysis included 6,418 protein-coding genes with a minimum expression of 50 reads per million in at least one sample. Clustering was performed using the Euclidean distance metric and average linkage. **(C)** Comparison of Venus transcripts levels between Venus-labeled DTCs and unlabeled samples across developmental stages. Transcript levels are shown as reads per million in log2 scale.

**A**

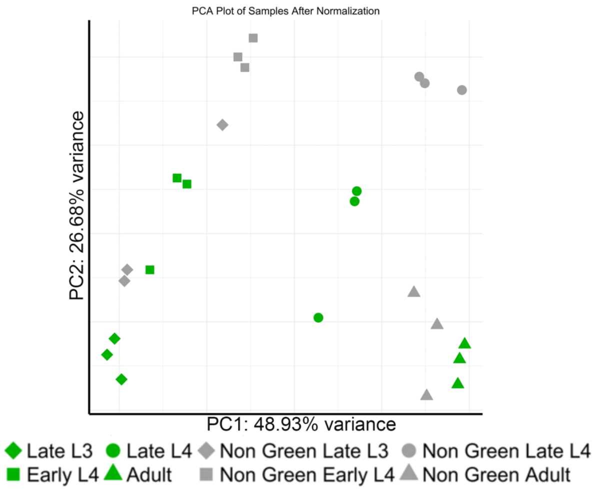


**B**

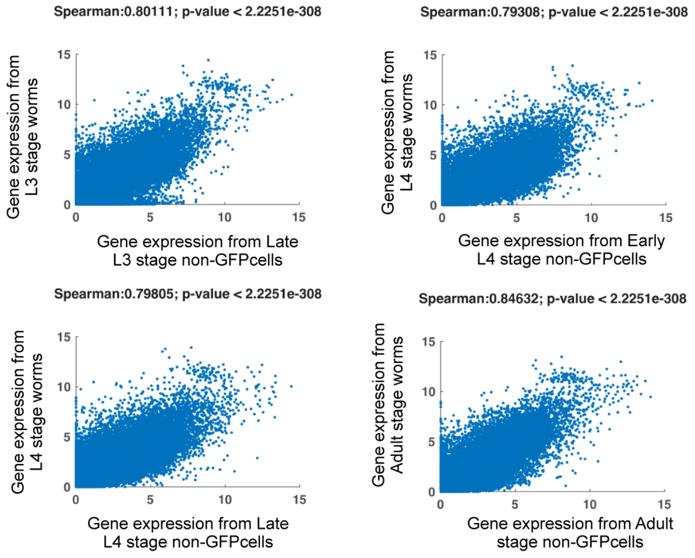


**Figure S2: Stage-specific expression of DTC transcripts.**

**(A)** Principal component analysis (PCA) of gene expression across Venus-positive (green; distal tip cells) and Venus-negative (gray; non-distal tip cells) samples from four developmental stages. Each point represents a replicate; developmental stages are indicated by symbol: diamonds for late L3, squares for early L4, circles for late L4, and triangles for adult. Percent variance explained by PC1 and PC2 is shown on the x- and y-axes, respectively.

**(B)** Comparison of averaged expression levels of protein-coding genes in non-Venus samples (X-axis) with median FPKM expression values from WormBase across a range of developmental stages (Y-axis). The WormBase dataset represents the median of all RNA-seq FPKM values for a given gene across wild-type samples at each developmental stage. Values are shown in log2 scale.

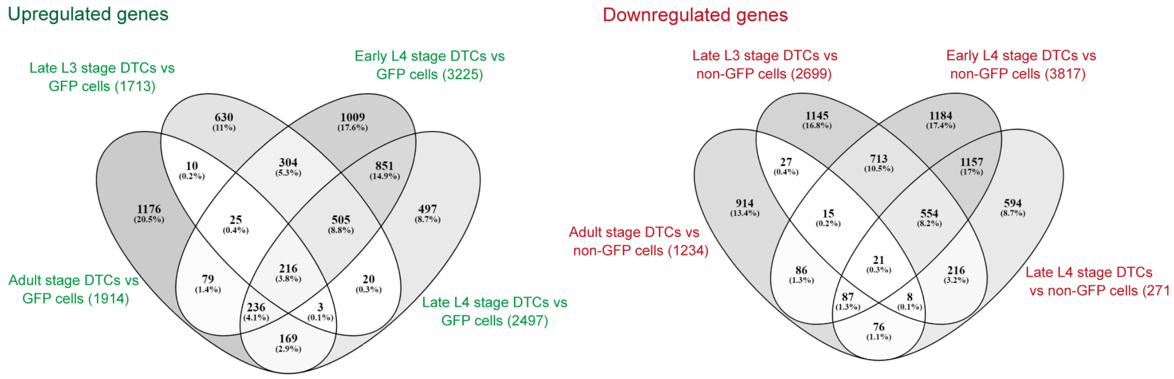


**Figure S3: Overlap of differentially expressed genes in DTCs between stages.** Venn diagrams depicting the overlap between genes upregulated (left) or downregulated (right) in late L3-, early or late L4- and Adult-stage DTCs.
