## Supplementary material for "Stage-specific transcriptomics of a leader cell reveals functional machineries driving collective invasion": Table S1

**Table S1: RNA sequencing alignment report**

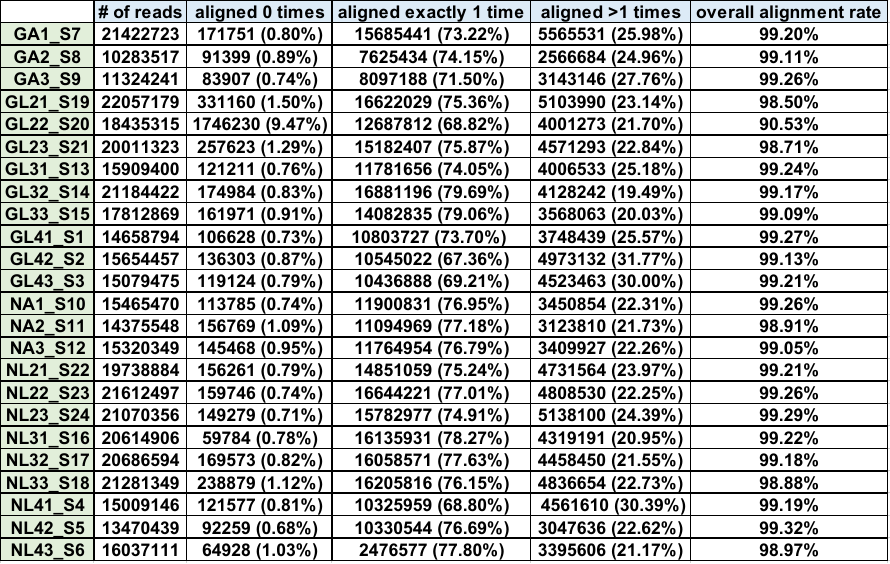

GA stands for Venus-positive adult stage DTC sample

GL2 stands for Venus-positive Late L3 stage DTC sample

GL3 stands for Venus-positive Early L4 stage DTC sample

GL4 stands for Venus-positive Late L4 stage DTC sample

NA stands for non-fluorescent adult stage DTC sample

NL2 stands for non-fluorescent Late L3 stage DTC sample

NL3 stands for non-fluorescent Early L4 stage DTC sample

NL4 stands for non-fluorescent Late L4 stage DTC sample
