## Supplementary material for "Stage-specific transcriptomics of a leader cell reveals functional machineries driving collective invasion": Table S4

**Table S4:** The genotype of *C. elegans* strains used in this study.

| **Strain** | **Genotype** | **Source** |
| --- | --- | --- |
| NF2169 | *mig-24p::Venus* | Gift from Kiyoji Nishiwaki |
| NK2115 | *cpIs121 I; rrf-3(pk1426) II; rde-1(ne219) V.* | 1 |
| RZB353 | *cpIs121 I; rrf-3(pk1426) II; rde-1(ne219) V;ltIs44 [Ppie-1::mCherry::PH(PLC1delta1); unc-119(+)] IV* | 2 |
| RZB431 | *muIs253[Peft-3::sfGFP_1–10_::unc-54 3'UTR Cbr-unc-119(+)] II; unc-119(ed3) III); sfGFP_11_:: sec-12* | This study |
| RZB425 | *muIs253[Peft-3::sfGFP_1–10_::unc-54 3'UTR Cbr-unc-119(+)] II; unc-119(ed3) III); rab-8::sfGFP_11_* | This study |
| RZB433 | *muIs253[Peft-3::sfGFP_1–10_::unc-54 3'UTR Cbr-unc-119(+)] II; unc-119(ed3) III); flwr-1::sfGFP_11_* | This study |
| NK2442 | *mig-6(qy37[mNG+loxP::mig-6]) V* | CGC |
| NK2413 | *sdn-1(qy29[sdn-1::mNG+loxP]) X* | CGC |
| LP362 | *gex-3(cp114[mNG-C1^3xFlag::gex-3]) IV. Hide Description* | CGC |
| LP439 | *nud-2(cp170[nud-2::mNG-C1^3xFlag]) I* | CGC |
| GLW16 | *rab-7(utx12[mNG::rab-7]) II* | CGC |
